# Phenotypic resistance to phage infection through capsule expression shutdown in *Klebsiella pneumoniae*

**DOI:** 10.1101/2025.06.16.659931

**Authors:** Lucas Mora-Quilis, Rafael Sanjuán, Pilar Domingo-Calap

**Affiliations:** Institute for Integrative Systems Biology, University of Valencia-CSIC, 46980 Paterna, Spain

## Abstract

Capsule loss is a major mechanism by which bacteria evade phage infection. This process has traditionally been attributed to mutations in capsule biosynthesis genes. Here, we investigated phage resistance in *Klebsiella pneumoniae*, a medically relevant encapsulated bacterium. Phage infection rapidly selected for resistant acapsular cells. As expected, transcriptomic analysis revealed a marked downregulation of capsule biosynthesis genes. However, full genome sequencing showed that capsule loss occurred without evidence of mutations, and acapsular phage-resistant cells were able to rapidly restore their capsule once phage pressure was removed. These findings highlight that phage-driven selective pressure can act on non-heritable variation in gene expression, providing a faster and more flexible resistance mechanism compared to the traditional mutation-selection process.

**IMPORTANCE:** Bacteriophages are increasingly considered as alternatives or complements to antibiotics, particularly against multidrug-resistant pathogens like *Klebsiella pneumoniae*. A key barrier to effective phage therapy is the rapid emergence of bacterial resistance. Capsule loss is a common resistance mechanism, traditionally linked to genetic mutations. Here, we show that *K. pneumoniae* can evade phage infection through reversible, non-mutational downregulation of capsule biosynthesis. This phenotypic adaptation allows rapid resistance development without compromising long-term fitness. Our findings reveal a flexible, non-genetic resistance strategy that may limit the durability of phage therapy and should be considered in future phage treatment designs.

## INTRODUCTION

Phages and their bacterial hosts are engaged in an arms race that has led to the evolution of diverse bacterial immune systems and phage countermeasures ^1–5^. When a naive bacterial population is challenged with a virulent phage, resistant bacteria usually emerge rapidly. The standard evolutionary process for resistance acquisition is the selection of mutants at loci encoding key infectivity factors such as phage receptors or other proteins involved in virus-host interactions ^6^. Bacterial diversity-creating mechanisms such as site-specific recombination and slipped-strand mispairing ^7,8^ can accelerate this process. However, it has long been reported that phage-sensitive bacteria persist in the presence of lytic phages or rapidly reappear upon phage clearance, questioning the standard evolutionary model and suggesting the involvement of additional and potentially non-genetic processes. These may include spatial refugia ^9^, herd immunity ^7^, but also variations in host physiology ^10^. For example, work with *Listeria* has shown that phage endolysins induce cell wall lesions leading to the formation of L-form wall-deficient cells resistant to phage-induced lysis ^11^. Spontaneous fluctuations in the expression levels of receptors and other entry factors can also lead to transient phage resistance ^12–14^. Rapid changes in host traits associated with resistance are often referred to as phase variation, but the underlying mechanisms of this process remain unclear in many cases. These may involve epigenetic modifications ^15^ or rapidly mutating loci ^7^.

Capsules play important roles in the virulence, environmental stability, and immune evasion capacity of some bacteria ^16^. Capsules are also important determinants of phage infectivity, as they can either protect cells from phages or be specifically used by phages for attachment and entry ^17–19^. Current wisdom states that capsule loss or capsular type shifts determined by sequence polymorphisms are common drivers of phage resistance ^20,21^. In *Bacteroides*, capsular type shifts can be driven by a genetic phase variation process involving the reorientation of genome fragments called recombinational shufflons, a mutational process that allows rapid responses to selection in capsule-determining loci ^22^.

An interesting model to study capsule-mediated phage-host interactions is *Klebsiella pneumoniae,* a clinically relevant pathogen exhibiting extensive capsular diversity, with more than 180 different capsular types described ^23^. Capsule biosynthesis in *K. pneumoniae* is mainly regulated by a Wzx/Wzy-dependent pathway, which is the most ubiquitous route for the production of bacterial polysaccharides with complex structures ^24–26^. The capsule operon consists of a core of genes encoding glycosyltransferases for the formation of the oligosaccharide chain, flippases for the translocation of the growing oligosaccharide across the inner membrane to the periplasm, and transporters for the export of the nascent polysaccharide across the outer membrane to the cell surface. Mutations in any of these genes can potentially affect capsule production. For example, point mutations and deletions can lead to acapsular mutants, as observed in the *rfaH* gene, blocking capsule formation ^27,28^. In addition, transposon-based libraries have shown that other genes may also be involved in capsule biosynthesis ^29^. Therefore, the genetic basis of *K. pneumoniae* capsule loss is well established.

Here, we aimed to characterize phage resistance via capsule loss in *K. pneumoniae*. We found that resistance appears rapidly, but that this process does not typically involve mutations in the bacterial genome. Instead, bacteria with downregulated capsule expression are selected in the presence of phage, and the capsular phenotype is restored rapidly after phage removal. This phenotypic resistance mechanism is faster, more versatile, and potentially more frequent than the standard mutation-selection evolutionary process.

## RESULTS

### Phage infection selects for capsule under-expression in the absence of mutations

*K. pneumoniae* wild-type cultures (WT) were challenged with phage Cap62, resulting in bacterial lysis. At 3 hours post inoculation (hpi), bacterial regrowth was observed in all replicates, indicating the emergence of phage-resistant bacteria (Figure 1A). The resulting resistant population (RP) exhibited an increased proportion of acapsular cells as shown by capsule staining and quantification (Figures 1B and 1C). As expected, no lysis was observed upon reinfection of RP with the phage (Figure 1B). To investigate the genetic basis of this process, WT and RP were sequenced but, surprisingly, no relevant mutations were found in RP despite the clear acapsular and resistant phenotype (Figure 2A and Data S1). All detected genetic variants were present at low population frequencies and showed no major deviations between WT and RP populations. To explore resistance mechanisms further, whole transcriptome sequencing was performed (Figure 2B). We found 19 genes showing a Log_2_FC > |3|, *p-adj < 0.00005*, of which 36.8% were up-regulated and 63.2% were down-regulated (Data S2). Notably, genes from the capsule biosynthesis locus such as *wcaI*, *wcaJ* and *gmd* were significantly down-regulated in the RP compared to the WT (Figure 2C), probably contributing to the acapsular phenotype. The operon responsible for anaerobic glycerol catabolism (*glpABC*) showed the most significant expression change (Figure 2D) (Log_2_FC < -3.96, *p-adj < 0.00005*).

**Figure 1.**
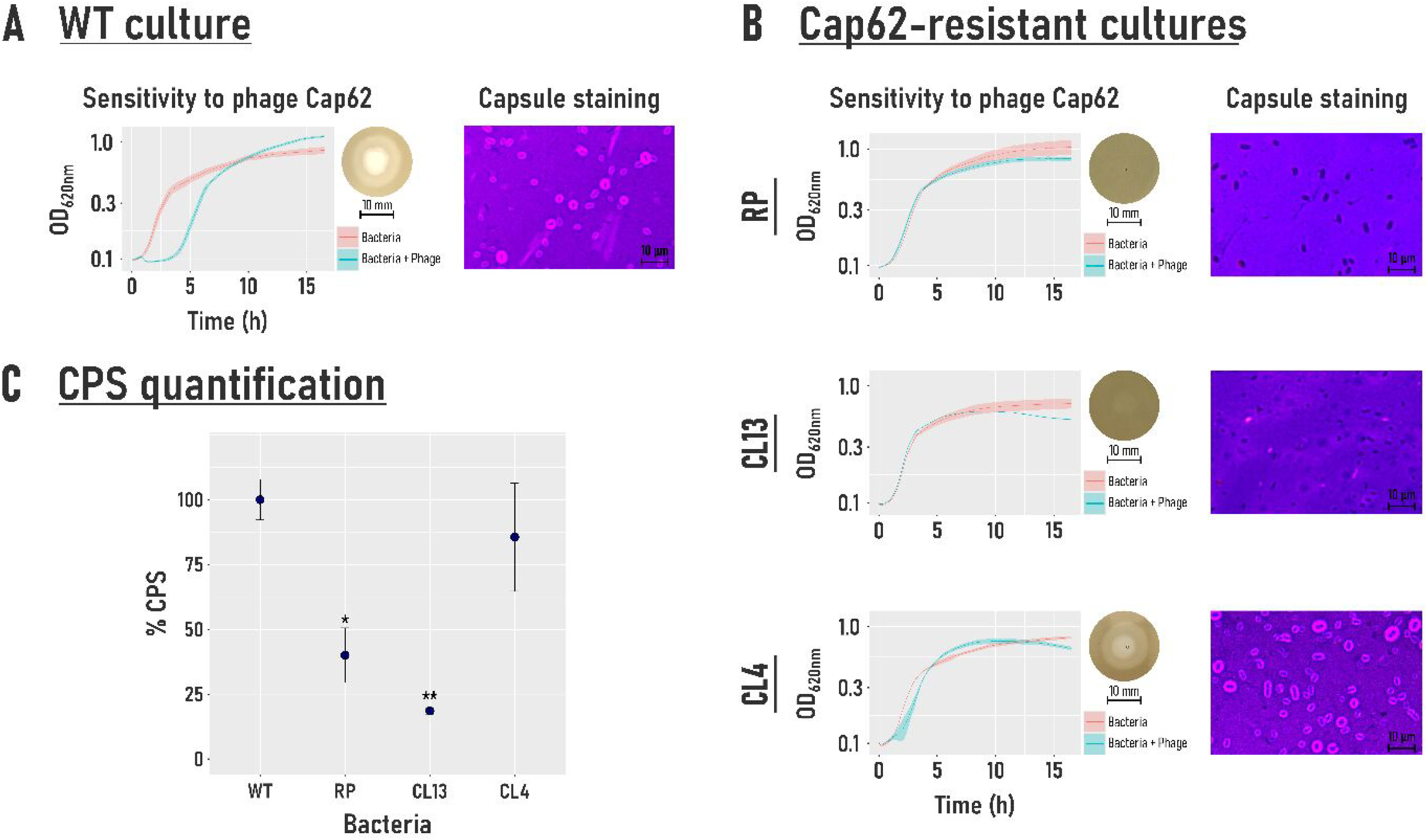
Phenotypic characterization of bacterial cultures. (A) WT culture. Growth curves of bacterial cultures (salmon line) and Cap62-infected cultures (blue line). OD_620 nm_ was measured over 16 hours, with three replicates per condition. Shaded areas represent the standard error of the mean (SEM). The microscopy image shows the capsule staining of WT culture. (B) Growth curves and capsule staining for Cap62-resistant cultures, including the resistant population (RP) and resistant clones CL13 and CL4 (*See Results section 3 for details*). The images reveal the presence or absence of capsule. (C) Capsule quantification relative to WT. Blue dots represent the mean for three replicates, and the error bars represent the SEM. \*\**p-value* < *0.001, *p-value* < *0.05, paired t-test*.

**Figure 2.**
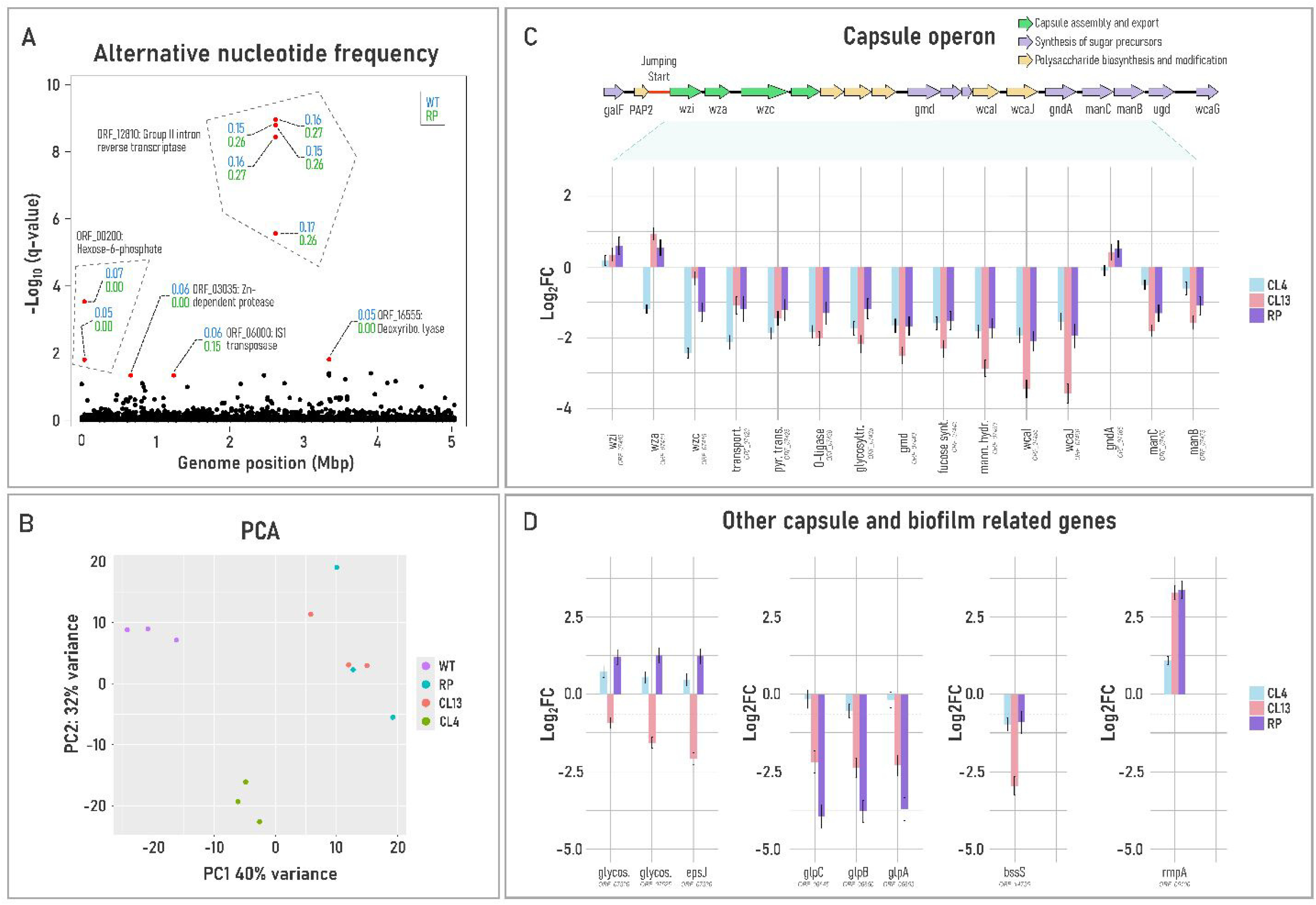
Whole genome sequencing and transcriptomic expression profile of Cap62-resistant cultures. (A) Alternative nucleotide frequency comparison between WT and RP across the chromosome. Each dot represents the q-value from a Fisher’s exact test, adjusted for a Discrete False Discovery Rate (FDR) test. Red dots indicate positions with q-values < 0.05 and an absolute difference in alternative allele frequency > 0.05. For each significant position, the corresponding gene name and open reading frame (ORF) are specified. Blue values represent the alternative nucleotide frequency in the WT, while green values represent those in the RP. Dotted grey lines group positions within the same gene. (B) Principal component analysis (PCA) of Cap62-resistant cultures. (C) Gene expression profile of genes included in the capsule biosynthetic cluster, relative to the WT culture, shown as Log_2_FC. Error bars indicate the log fold change standard error. Expression values below the dotted grey line are not statistically significant (*p-adj > 0.05*, Wald test). (D) Gene expression of other operons and genes related to capsule and biofilm biosynthesis. Error bars indicate the log fold change standard error. Expression values below the dotted grey line are not statistically significant (*p-adj > 0.05*, Wald test).

### Capsule synthesis and phage sensitivity are rapidly restored after phage removal

To better characterize this process, individual cells from the WT or RP were isolated by flow cytometry and incubated in separate wells in the presence or absence of phage. At 48 hpi, the number of wells with bacterial growth was determined to infer the proportion of phage-resistant cells present in WT and RP (Figure 3A). Whereas WT was fully composed of sensitive cells (0% growth in presence of phage), 92% of the RP cells were found to be resistant, confirming the above observations. Then, individual cells from RP were grown either in the absence or presence of phage (Figure 3B). Capsule staining revealed that cultures regrown in the absence of phage rapidly restored their capsule, whereas cultures regrown in the presence of phage remained acapsular. These cultures were again subjected to cell-sorting to determine the percentage of resistant cells, which decreased to 66% after the first passage in the absence of phage, and to 51% after a second passage. In contrast, when passages were performed in the presence of phage, the proportion of resistant cells remained above 90%.

**Figure 3.**
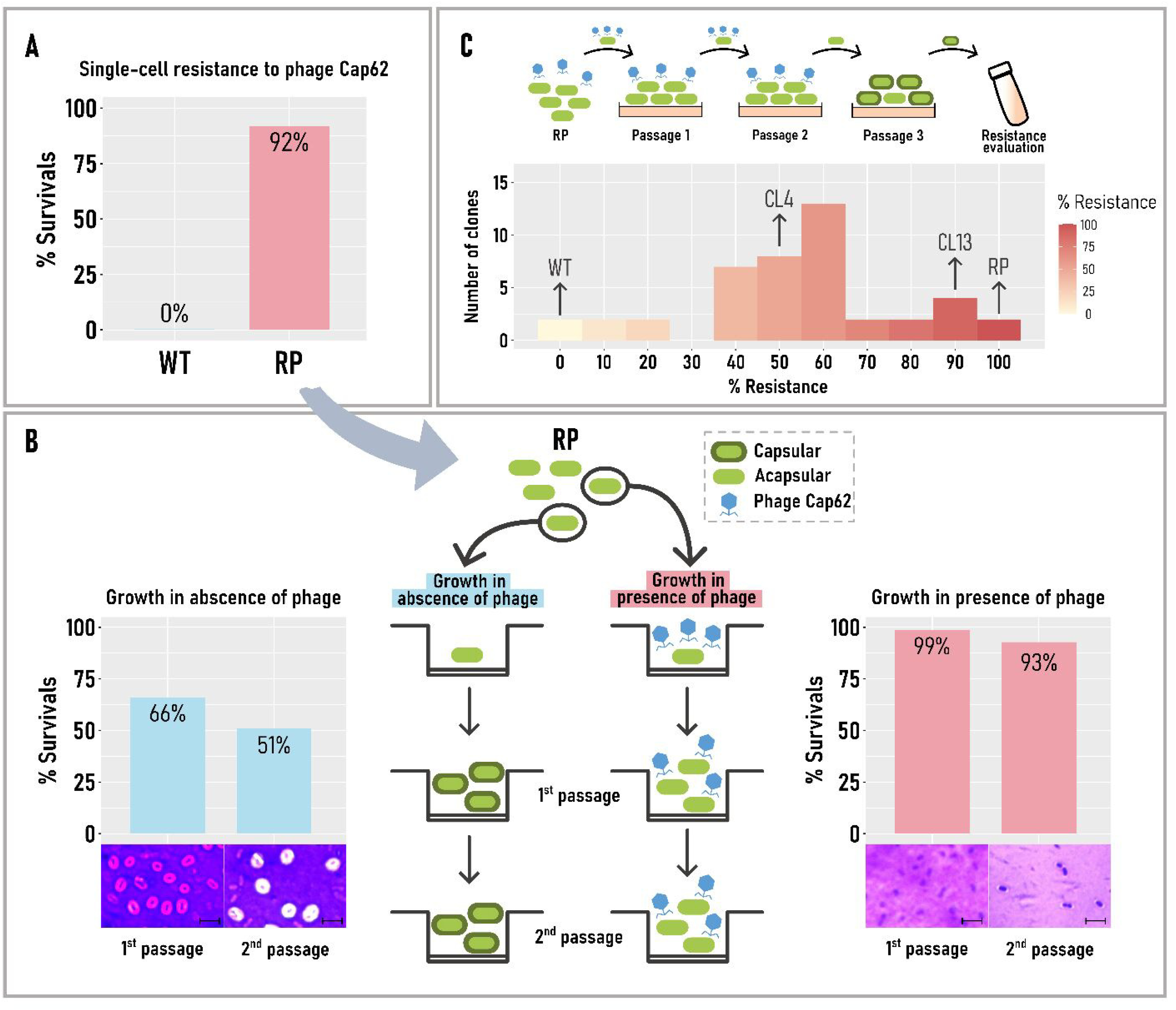
Capsule synthesis dynamics in the presence and absence of phage selective pressure. (A) Proportion of Cap62 resistant cells in WT (blue) and RP (salmon) cultures. (B) Acapsular resistant cells from the RP were isolated by single-cell sorting and grown in the absence (blue) or presence (salmon) of phage Cap62. Acapsular cells grown in the absence of phage restored capsule production, increasing the proportion of cells sensitive to the phage. In contrast, acapsular cells grown in the presence of phage remained acapsular and highly resistant. Scale bars in microscopy images represent 5 µm. (C) Illustration of colony isolation and phenotypic change: Acapsular cells from the RP were streaked onto agar plates to select a single cell-derived colony (passage 1). The acapsular phenotype was initially maintained by residual phage in the environment. Colony passage was repeated at least twice (passages 2 and 3) until phage Cap62 was completely removed. Note that the removal of Cap62 allowed capsule restoration. Finally, each individual colony was grown and tested for resistance to Cap62. The histogram shows the resistance distribution of 42 selected colonies based on both spot and liquid infections. WT and RP cultures represent the minimum and maximum resistance levels, respectively.

### Mutations and gene expression changes provide alternative mechanisms of resistance

To further examine capsule restoration after phage removal, 43 colonies were isolated from RP and grown independently. Each of these clones was then challenged with phage to assess resistance levels. The results showed variable outcomes, ranging from complete resistance to sensitivity levels similar to the WT (2.99% - 95.52% resistance; Figure 3C and Data S3). Clone CL4 was selected as a representative of intermediate resistance, and CL13 as representative of stable resistance. Whereas capsule production was restored in CL4, CL13 remained acapsular (Figure 1B). Genomics and whole transcriptomics were performed for the two clones to compare them with WT and RP (Figure 2). No mutations were found in CL4, as previously observed in RP. However, CL13 had a 3 nucleotide in-frame deletion in the *rfaH* gene, resulting in a valine deletion. RfaH is a transcriptional elongation factor involved in the biosynthesis of capsular polysaccharides, lipopolysaccharides and other virulence factors. The deleted valine was located at position 26 in the N-terminal domain of the protein, which is involved in the interaction with the operon polarity suppressor, a transcriptional pausing signal.

Transcriptomics revealed a general down-regulation of the capsule biosynthesis genes cluster in both CL4 and CL13 cultures compared to WT (Figure 2C). In CL13, gene expression levels decreased with distance from the start of the operon, indicating a dysfunction of the RfaH transcription elongation factor due to the valine deletion. The expression levels of genes such as *wcaI*, *wcaJ* and *CDS_07450* were significantly reduced (Log_2_FC < -2.5, *p-adj < 0.00001)*, which may contribute to the acapsular phenotype. Additionally, the *epsJ* and the *bssS* genes, related with the synthesis of capsule and biofilm polysaccharides, respectively, were down-regulated (Log_2_FC < -2, *p-adj < 0.00001*) (Figure 2D). Interestingly, the mucoid phenotype regulator *rmpA*, was highly over-expressed in the acaspular cultures, potentially as a compensatory response to the lack of capsule (Log_2_FC > 3.3, *p-adj <0.00001*). However, in CL4 and RP, the expression levels of the capsule operon remained relatively low despite the absence of mutations. Surprisingly, CL4 showed significantly lower expression levels for certain genes, such as *wza* and *wzc*, compared to the RP (Log_2_FC < -1.1, *p-adj < 0.001*) (Figure 2C). The only gene with significantly lower expression in RP compared to CL4 was *manC* (Log_2_FC < -0.8*, p-adj < 0.0005*). However, these differences are unlikely to be sufficient to explain the acapsular phenotype in RP compared to CL4. Notably, the *glpABC* operon, which was greatly downregulated in RP, exhibited expression levels in CL4 similar to those in WT (Log_2_FC < - 0.38, *p-adj < 0.05*) (Figure 2D). Therefore, the downregulation of these genes may be indirectly related to the acapsular phenotype observed in RP.

## DISCUSSION

The mechanisms by which capsule synthesis is lost in response to phage infection remains to be elucidated. Capsule production is governed by complex gene expression networks involving multiple regulators and environmental signals ^29^. One possibility is that signalling molecules secreted by infected cells could trigger this downregulation in uninfected cells, making them resistant to phage entry. We could not find evidence for such signalling, since filtered supernatants from infected cultures did not modify capsule production in naïve bacteria. We therefore suggest that capsule loss was driven by selection of spontaneous gene expression variants already present in the population before phage infection. This was probably the most prevalent resistance emergence mechanism in our experiments, as no significant change in allele frequencies was associated with capsule loss and phage resistance. Most of the selected variants rapidly restored capsule validating the non-genetic regulation of capsule production, and only a small fraction of the population retained a stable acapsular phenotype due to mutations. Our results are consistent with previous observations in *Salmonella* spp., where epigenetic regulation of the *opvAB* operon produces a shorter O-antigen, reducing virulence but enhancing phage resistance ^15^. Notably, phase variation allows rapid restoration of normal LPS variants once phage pressure subsides ^14^, as observed here for capsule production. In *K. pneumoniae*, the down-regulation of genes such as *wzc*, *gmd*, *wcaI*, and *wcaJ* has been previously associated with the acapsular phenotype, in agreement with our results ^29–31^.

Notably, the operon responsible for anaerobic glycerol catabolism (*glpABC*) showed the most significant expression change in response to phage infection. The downregulation of these genes leads to a general energy shutdown linked to increased persister cell formation. Persister cells are characterized by a dormant, low-energy state that enhances their tolerance to certain antibiotics ^32–35^. This metabolic adaptation likely reduces energy production and the biosynthesis of energy-intensive structures, including the capsule and other surface components ^36–38^. Mutations in *glpC* have been shown to impair biofilm formation ^39^. These findings suggest that stress conditions, such as phage infection, may drive the down-regulation of *glp* genes, promoting a low-energy metabolic state that indirectly selects for acapsular, phage-resistant variants.

The evolutionary implications of phase variation driven by changes in gene expression in the absence of mutations need to be further investigated. Phenotypic plasticity is a form of robustness that promotes survival in changing environments and hence affords additional time for the emergence and selection of beneficial mutations, but robustness also reduces fitness differences between genetic variants, reducing the efficiency of selection. ^40,41^.

Future work may elucidate whether similar mechanisms of adaptation occur in other encapsulated bacteria, such as *Acinetobacter baumannii, E. coli* or *Salmonella* and in response to other selective pressures, such as host immunity or antibiotic treatments. This would lead to new models of bacterial evolution and provide valuable insights for the development of therapeutic strategies.

## MATERIALS AND METHODS

### Bacterial strain and phage

The bacteria used in this study was *K. pneumoniae* capsular type-1, a reference strain from the collection of the Statens Serum Institute (Copenhagen, Denmark). Bacteria were grown in lysogeny broth (LB) supplemented with 3.78 mM CaCl_2_ at 37°C with shaking. Wastewater samples were used for phage isolation as described previously ^42^. The phage used in this study was *Klebsiella* phage vB_Kpn_Cap62 (Cap62) (PQ741857), isolated using K1 as the primary host. Phage Cap62 belongs to the family *Autographiviridae* and the genus *Drulisvirus*. The genome size is 43906 base-pairs (bp) and it encodes a receptor-binding protein with depolymerase activity targeting the K1 capsule, as described for a closely related phage ^43^.

### Selection and isolation of phage-resistant bacteria

Wild-type (WT) was inoculated with phage Cap62 until emergence of resistant variants. The resulting culture was designated as the resistant population (RP). To isolate phage-free resistant clones, the resistant cultures were washed three times and streaked onto LB agar plates. At least three sequential colony-to-colony passages were performed to eliminate the residual phage. Individual colonies were then picked, grown in LB to stationary phase, and the supernatants were tested to confirm the complete absence of phage. These phage-resistant cultures were subsequently stored in 20% glycerol at -70°C.

### Capsule staining and quantification of bacterial cultures

To determine the presence of the capsule in bacterial cultures, contrast staining with crystal violet and nigrosine was performed ^42^. Liquid cultures with a minimum recommended concentration of 10^9^ colony forming units (CFU)/mL were fixed for 20 min with 2.5% formaldehyde in the presence of 100 mM lysine. The fixative agent was removed by centrifugation and washed with phosphate-buffered saline (PBS). 25 – 40 µL of the fixed culture was mixed with a small drop of nigrosine 10% in a glass slide and smeared along the slide until air dried. 1% crystal violet was gently poured to cover the slide and incubated for 5 min at room temperature. The preparation was carefully rinsed with distilled water, air dried and examined under a light microscope. Capsule production was quantified using the uronic acid method ^19^. Bacterial cultures were washed twice with PBS. For capsule extraction, 500 µL of the bacterial suspension was mixed with 100 µL of 500 mM citric acid and 1% Zwittergent 3-10. After a 20 min incubation at 56°C, cellular debris was pelleted, and the supernatant was mixed with ethanol (final concentration 80% v/v) and incubated for 30 min at 4°C to precipitate the capsule. Tubes were centrifuged (14000 ×g, 20 min, 4°C), air-dried, resuspended in water, and incubated for 2 h at 56°C. To quantify capsule, 1.2 mL of borax solution was added to the precipitated capsule and incubated at -4°C for 10 min, followed by a 10-min incubation at 95°C. Afterward, the samples were incubated at -4°C for 10 min. Finally, 20 µL of 3-hydroxybiphenyl in 0.5% NaOH was added as a colorimetric agent to quantify the uronic acid present in each sample, using a standard curve with glucuronic acid for comparison.

### Evaluation of bacterial phage-resistance

The resistance of bacterial cultures to phage Cap62 was assessed by the spot-test technique and liquid infections. For spot-test, 200 µL of a stationary phase culture containing approximately 10^9^ CFU was mixed with 3.5 mL of semi-solid agar LB and plated it onto an LB plate. After drying, 1 µL of phage was spotted at a concentration of approximately 10^8^ plaque-forming units (PFU)/mL. In addition, spots of ten-fold serial dilutions were also routinely performed. After an over-night (ON) incubation at 37°C clear spots were scored as 2, and turbid spots as 1 and the absence of spot as 0. Each spot was evaluated spot was evaluated in at least triplicate in independent experiments. For liquid infections, approximately 5 × 10^6^ CFU were mixed with phage to the desired concentration in a final volume of 150 µL. To track phage lysis and the growth of the resistant bacteria, we measured the OD_620_ nm over time with background shaking at 37°C, using 96-well plates and a plate reader (Multiskan FC). To determine the resistance in liquid infections, the slope for each two adjacent points was calculated as 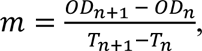 where *OD* represents the OD_620_ nm, and *T* is the time. The time to resistance was determined by calculating the difference between the time points at which the maximum slope was reached for untreated bacteria and bacteria treated with phages. All experiments were performed in triplicate.

### Single-cell resistance to phages using cell-sorting

Bacterial cultures were grown in LB medium or in presence of phage. To remove the broth culture and partially eliminate the surrounding phage, cultures were washed five times with PBS. These were then resuspended in PBS to a final concentration of 10^5^ CFU/mL. These suspensions were subsequently passed through a cytometer (BD FACSAria™ Fusion) under biosafety level-2 conditions to isolate individual cells into 96-well plates containing either LB or phage Cap62 at a minimum concentration of 10^8^ PFU/mL, ensuring that only resistant bacteria could grow. Plates were incubated for 48 h at 37°C, and the number of wells with bacterial growth was counted. The number of wells with bacterial growth in LB medium represented the maximum growth (100%). The proportion of wells with bacterial growth in the presence of phage was then normalized to this maximum growth to determine the relative resistance of the bacteria to the phages.

### Conditioned medium assay

To investigate whether acapsularity in *K. pneumoniae* could be induced by signalling molecules secreted by phage-resistant bacteria in the presence of phage, resistant cells were removed from RP cultures by centrifugation (10000 ×g, 3 min, 4°C). The supernatants were filtered twice through a 0.05 µm filter to ensure complete removal of phage Cap62 while retaining any potential signalling molecules. Fresh K1WT bacteria were inoculated into these conditioned supernatants and incubated at 37°C until reaching the late exponential growth phase. The presence of capsule presence was subsequently assessed microscopically, following the previously protocol.

### Bacterial DNA sequencing and genomic analysis

Bacterial cultures were grown to stationary phase and DNA was extracted using the PureLink Genomic DNA Mini Kit (Invitrogen). Free-PCR 150 paired-end libraries were prepared using the NEBNext Ultra IIDNA Library Prep Kit, and sequenced using the Illumina NovaSeq6000 platform. Read quality was assessed using FastQC ^44^, and genomes were assembled against a reference genome (CABDWB000000000) using bwa-mem2 ^45^ to align the reads. To analyse single nucleotide variants (SNVs), Snippy ^46^ and LoFreq ^47^ were used with default parameters. BreakDancer Max was used to detect large genomic variants ^48^. Reads were mapped to the wild-type genome, and variants were manually inspected using IGV ^49^. In addition, to discard false positives predicted by the previous variant callers, we extracted the nucleotide counts for each allele across the entire genome and applied a False Discovery Rate (FDR) analysis for discrete data to compare these counts between WT and RP cultures.

### Bacterial mRNA sequencing and transcriptomic analysis

Total RNA from exponential phase cultures (in triplicate) was extracted using RNAzol RT (Sigma-Aldrich) according to the manufacturer’s protocol. RNA concentration was measured with the Qubit Fluorometer (Thermo Scientific), and RNA quality and integrity was assessed using Nanodrop (Thermo Scientific) and the 2100 Bioanalyzer (Agilent). RNA ribosomal depletion was performed prior to cDNA library construction, and sequencing was carried out using the Illumina NextSeq 500 platform (2 ͯ 75 bp). Transcripts were mapped to the WT genome using HISAT2 ^50^, and total read counts for each genomic feature were extracted with featureCounts ^51^. Differential transcription analysis between samples was performed using DESeq2 ^52^ with read counts as input.

### Quantification and statistical analysis

To estimate the differences in capsule quantification between samples, paired t-tests were performed on measurements in triplicates using the R software (v4.3.3) ^53,54^. Significant differences in allele counts at each genomic position were estimated using Fisher’s exact test, with FDR correction applied via the Discrete Benjamini-Hochberg method, implemented in the DiscreteFDR package (v2.0.0) ^55^ in R. Significant differences in transcription levels between WT and resistant cultures were analyzed using DESeq2 (v1.42.1) ^52^ in R, with normalization performed via the variance-stabilizing transformation function (vst). Statistical significance of Log_2_FC was assessed using the Wald test. Principal component analysis (PCA) was conducted in R to visualize gene expression variance between samples, while relative gene expression to WT was plotted as bar graphs for selected operons using ggplot2 (v3.5.1) ^56^.

## ACKNOWLEDGMENT

This research was funded by project PID2020-112835RA-I00 funded by MCIN/AEI /10.13039/501100011033, and project SEJIGENT/2021/014 funded by Conselleria d’Innovació, Universitats, Ciència i Societat Digital (Generalitat Valenciana) to P.D-C. P.D-C. was financially supported by a Ramón y Cajal contract RYC2019-028015-I funded by MCIN/AEI/10.13039/501100011033, ESF Invest in your future. L.M-Q. was funded by a PhD fellowship FPU19/04611 from Spanish MCIU.

## SUPPLEMENTAL INFORMATION

### Data S1. Alternative allele frequency differences between WT and RP cultures

For simplicity, only the 1000 genomic positions with the most significant difference in alternative allele frequency are shown. For each position, the consensus (cons) and alternative (alt) allele counts are provided for both WT and the RP cultures. The alternative allele frequency (AF) is calculated as the ratio of alternative allele counts to total allele counts in each culture. The absolute difference in alternative allele frequency (|ΔAF|) is defined as |AF_RP – AF_WT|. Positions with q-value < 0.05 (Fisher’s exact test with FDR correction) and |ΔAF| ≥ 0.05 are highlighted, with gene features specified.

### Data S2. Gene expression in phage-resistant cultures relative to WT

For simplicity, only genes with p-adj < 0.0001 are shown. Data for each phage-resistant culture (RP, CL13 and CL4) is presented in separate Excel sheets. For each culture, the table includes open reading frames (ORF), Log_2_FoldChange (L2FC) relative to WT expression, adjusted p-value determined by the Wald test (p-adj), regulation status (up- or down-regulated), and other features of interest such as sequence type, gene name and product name.

### Data S3. Resistance profile of phage-resistant cultures

Resistance of 46 phage-resistant clones is shown in comparison to WT (lowest resistance) and the RP (highest resistance). Resistance was assessed using spot and liquid infection assays. Spot sensitivity was scored as 2 (clear lytic spots), 1 (turbid spots), and 0 (no lytic spots). Liquid sensitivity was measured as the time to emergence of resistance based on OD_620_ nm with the standard error of the mean (SEM). Relative resistance to WT (%) was calculated by integrating the data from both assays.

